# Prediction of sex-determination mechanisms in avian primordial germ cells using RNA-seq analysis

**DOI:** 10.1101/2022.02.24.481709

**Authors:** Kennosuke Ichikawa, Yoshiaki Nakamura, Hidemasa Bono, Ryo Ezaki, Mei Matsuzaki, Hiroyuki Horiuchi

## Abstract

Avian sex is determined by various factors, such as the dosage of DMRT1 and cell-autonomous mechanisms. While the sex-determination mechanism in gonads is well analyzed, the mechanism in germ cells remains unclear. In this study, we explored the gene expression profiles of male and female primordial germ cells (PGCs) during embryogenesis in chickens to predict the mechanism of sex-determination. Male and female PGCs were isolated from blood and gonads with a purity > 96% using flow cytometry and analyzed using RNA-seq. Prior to settlement in the gonads, female circulating PGCs (cPGCs) obtained from blood displayed sex-biased expression. Gonadal PGCs (gPGCs) also displayed sex-biased expression, and the number of female-biased genes detected was higher than male-biased genes. The female-biased genes in gPGCs were enriched in some metabolic processes. To reveal the mechanisms underlying the transcriptional regulation of female-biased genes in gPGCs, we performed stimulation tests. Stimulation with retinoic acid against cultured gPGCs derived from male embryos resulted in the upregulation of several female-biased genes. Overall, our results suggest that sex determination of avian PGCs possess aspects of both cell-autonomous and somatic cell regulation. Moreover, it appears that sex determination occurs earlier in females than in males.

## Introduction

Birds have unique mechanisms of sex-determination. Mammals, which have an XX/XY sex chromosome system, determine their gonadal sex depending on the transient action of a Y-chromosome linked master gene, *Sex-determining region Y* (*SRY*) (Koopman et al., 1991). In the case of birds (ZZ/ZW), it is likely that *Doublesex and mab-3 related transcription factor 1* (*DMRT1*) on the Z chromosome act as a dose-dependent regulator of gonadal sex differentiation (Smith et al., 2009a; Lambeth et al., 2014; Ioannidis et al., 2021). Expression of *DMRT1* is restricted to the developing gonads of both sexes but is more highly expressed in males than in females due to the Z-linked gene. Avian somatic cells possess an inherent sex identity, and gonadal development and sexual phenotype is largely cell autonomous (Zhao et al., 2010). Nevertheless, sex hormones also principally regulate sexual phenotypes, and thus sex-reversal, a characteristic feature in fish, is also observed in birds. Investigation of the mechanism in birds will elucidate evolution in vertebrate sex-determining mechanisms.

Little is known about the sex-determining mechanism of avian germ cells compared to that of somatic cells. The molecular mechanisms of sex determination in germ cells generally have received much attention in the past decade. In vertebrates, *Forkhead box L3* (*FOXL3*) was the first identified gene associated with the sex determination of germ cells in medaka (*Oryzias latipes*) (Nishimura et al., 2015; Tanaka, 2016). Development of functional sperm in the ovary of *FOXL3* knockout female medaka, suggested that the germline has a sex-determination mechanism different from that of gonadal sex in at least some vertebrates. However, in chickens, the *FOXL3*-like gene is temporally expressed in oogonia, which develop after the sex-determination of germ cells (Ichikawa et al., 2019). Therefore, the sperm-egg fate decision via *FOXL3* was not conserved in chickens. To understand the comprehensive mechanism for avian sex determination, avian-specific sperm-egg fate determinants must be revealed.

To elucidate the sex-determination mechanism of avian germ cells, gene profiling in primordial germ cells (PGCs), the first germ cell population established during early development, obtained from various developmental stages, is required. Sex-differentiation of germ cells is generally induced by the surrounding gonadal somatic cells. Since gonadal masculinization related genes, *DMRT1* and *Hemogen* (*HEMGN*), or feminization-related genes, *Forkhead box L2* (*FOXL2*) and *aromatase*, begin to be expressed in chicken embryos incubated for 4.5-5.7 days (E4.5-5.7) (Govoroun et al., 2004; Nakata et al., 2013; Lambeth et al., 2014), the sex-differentiation of gonadal PGCs (gPGCs) might also be induced from this stage onwards. Interestingly, cell-autonomous sex determination of chicken PGCs was also observed before the cells settled into the gonads. Unlike other vertebrates, avian PGCs use blood circulation for transport to the genital ridge. Although the precursor cells for PGCs transplanted to the opposite sex of chicken embryos at the blastodermal stage could differentiate into functional gametes (Kagami et al., 1997), circulating PGCs (cPGCs) transplanted into the bloodstream do not differentiate into functional gametes (Naito et al., 1999; Tagami et al., 2007). These previous studies suggested that the sex of PGCs is independent of gonadal differentiation. Therefore, to reveal the sex-determination mechanisms in avian PGCs, the effects of both surrounding gonadal cells and cell-autonomous factors must be considered.

In order to investigate the sex-determination mechanisms, we purified male and female PGCs from blood (E2.5; Hamburger and Hamilton stage (HH) 17) (Hamburger and Hamilton, 1951) and gonads (E4.5 and E6.5; HH25-26 and HH30, respectively) using fluorescence-activated cell sorting (FACS). Gene expression profiles of PGCs at each developmental stage for each sex were determined using RNA-seq analysis. Then, the sex-determination mechanism of PGCs was predicted by bioinformatic analysis. To evaluate the prediction, male PGCs were stimulated with retinoic acid *in vitro,* and the changes in gene expression were examined.

## Materials and methods

### Experimental animals

Fertilized eggs of White Leghorn were purchased from Akita foods (Fukuyama, Japan) and used for PGC collection from blood and gonads. All animal care and use in this study were conducted in accordance with the animal experimentation guidelines of the Hiroshima University Animal Research Committee.

### Sample preparation using flow cytometry

Male and female PGCs were isolated from the blood of E2.5 embryos and from the gonads of E4.5 and E6.5, embryos using FACS. Fertilized eggs were incubated at 37°C under 50-60% relative humidity. Collected gonad cells were dissociated using TrypLE™ Express Enzyme (Thermo Fisher Scientific, Carlsbad, CA, USA). The gonad cells and blood cells were incubated with anti-stage-specific embryonic antigen-1 (SSEA-1) antibody (sc-2170; Santa Cruz Biotechnology, Dallas, TX, USA) diluted with wash buffer (0.5% BSA and 0.1% NaN3-PBS) at 1:100 for 1 h on ice. After washing with the washing buffer, the secondary antibody reaction was carried out using FITC Rat anti-mouse IgM antibody (553408; BD Biosciences, Franklin Lakes, NJ, USA) at 1:200 dilution for 30 min on ice. After washing, the samples were incubated with propidium iodide to detect the dead cells. Then the SSEA-1 positive cells were sorted using Cell Sorter MA900 (Sony Biotechnology, San Jose, CA, USA). The sex of each embryo was confirmed by the patterns of *chromatin-helicase DNA-binding protein-1* (CHD1) fragment bands obtained by PCR (Ellegren, 1996; Griffiths et al., 1998). Genomic DNA was purified from a small section of embryos using SimplePrep™ reagent for DNA (TaKaRa Bio, Kusatsu, Japan).

### Immunofluorescence staining

The purity of the sorted cells was confirmed by immunofluorescence staining using an anti-chicken VASA homologue (CVH) mouse monoclonal antibody (Nakano et al., 2011). First, the sorted cells were adhered to slides with Smear Gell (GenoStaff, Tokyo, Japan) according to the manufacturer’s instructions. Subsequently, the adhered cells were fixed with 4% paraformaldehyde in PBS for 30 min at room temperature. After washing with 0.1% BSA-PBS, the adhered cells were incubated with 0.1% Tween-20. Subsequently, they were blocked with 3% BSA, and then the primary antibody reaction using hybridoma supernatant containing the anti-CVH antibody, which was diluted 1:1 with 2% BSA-PBS, was performed for 40 min at 37°C. Next, the cells were washed, and the secondary antibody reaction was performed using goat anti-mouse IgG (H+L) (Alexa Fluor Plus 555; Invitrogen, Waltham, MA, USA) at 1:200 dilution with 1% BSA-PBS for 40 min at 37°C. Finally, nuclei were stained with 4’,6-diamidino-2-phenylindole (DAPI) in VECTASHIELD Mounting Medium (Vector Laboratories Inc., Burlingame, CA, USA) after washing. They were observed under a fluorescence microscope (BX53; Olympus, Tokyo, Japan) and photographed with a DP74 camera (Olympus).

Furthermore, cultured male PGCs obtained from the gonads of E6.5, were centrifuged and fixed with 4% paraformaldehyde in PBS for 30 min at room temperature after washing. Immunofluorescence staining was performed using the same procedure as described above until the primary antibody reaction. After washing, the secondary antibody reaction was carried out with goat anti-mouse IgG (H+L) (Alexa Fluor Plus 488; Invitrogen) under the same conditions as described above. They were then washed and mounted on glass slides with VECTASHIELD Mounting Medium and observed under a fluorescence microscope.

### RNA-seq and bioinformatic analyses

Two pools of sorted PGCs from to 6-14 individuals of each sex and stage were used as templates. Since the yield of total RNA was very low, cDNA libraries were prepared using SMART-seq v4 Ultra Low Input RNA Kit for Sequencing (TaKaRa Bio), which is based on the SMART-seq2 method (Picelli et al., 2013). Then Nextera XT DNA Library Preparation Kit (Illumina Inc., San Diego, CA, USA) was used to make cDNA libraries suitable for Illumina sequencing. These were analyzed using the Hiseq system (Illumina Inc.) with 150 bp paired-end sequencing.

The quality of the sequencing results was assessed using FastQC (ver. 0.11.9). Trimming and quality filtering of the raw reads were performed using Trimmomatic (ver. 0.39) (Bolger et al., 2014) as well as Trim Galore (ver. 0.6.6) with Cutadapt (ver. 3.1) (Marcel, 2011) software. The trimmed and filtered reads were mapped to the chicken reference genome (GRCg6a) using STAR (ver. 2.7.3a) (Dobin et al., 2013). The uniquely mapped reads in each gene were counted using the featureCounts tool (ver. 2.0.1) (Liao et al., 2014). While detection of sex-biased genes was performed using the DE-seq2 tool (ver. 1.20.0) (Love et al., 2014). MA-plots were created using the Bokeh visualization library (ver. 2.3.2). Then, gene ontology (GO) analysis and Kyoto Encyclopedia of Genes and Genomes (KEGG) pathway analyses were performed using the g:Profiler online tool (Raudvere et al., 2019).Finally, tissue or cell specificity was characterized using the Metascape online tool (Zhou et al., 2019).

### RT-qPCR and RT-PCR

To confirm the results of RNA-seq, RT-qPCR analysis was conducted. In this analysis, whole transcriptome amplification was performed using the QuantiTect Whole Transcriptome Kit (QIAGEN, Venlo, Netherlands) against total RNA from new pools of sorted PGCs from 45-70 to embryos at each stage and sex. To remove any bias in the whole transcriptome amplification reaction, the reaction was performed in triplicate for each stage and sex (n = 3). The synthesized cDNA was then used for RT-qPCR using a StepOne real-time PCR system (Applied Biosystems, Waltham, MA, USA) with the KOD SYBR qPCR Mix (Toyobo Co. Ltd., Osaka, Japan). For the stable PCR reaction, a total of 0-10% concentration dimethyl sulfoxide was used. The primers used for this analysis are listed in Table S1. The RT-qPCR conditions were 40 cycles of 98°C for 10 sec, 60-65°C for 10 s, and 68°C for 30 s. Melting curve analysis was performed following this amplification stage in three steps: 95°C for 15 s, 60°C for 1 min, and 95°C for 15 sec. Gene expression analysis of cultured PGCs stimulated with retinoic acid (RA) was also conducted using RT-qPCR. Total RNA was purified using the RNeasy Micro Kit (QIAGEN). For cDNA synthesis, SuperScript IV reverse transcriptase (Thermo Fisher Scientific) was used against 100 ng of total RNA. The RT-qPCR reaction was also performed using the StepOne real-time PCR system (Applied Biosystems) with the KOD SYBR qPCR Mix (Toyobo Co. Ltd.), and the primers used in this reaction are shown in Table S2. The RT-qPCR conditions were as follows:40 cycles of 98°C for 10 sec, 60-62°C for 10 s, and 68°C for 30 s, followed by melting curve analysis under the same conditions described above.

Relative expression scores were calculated using the ΔΔCt method (Livak and Schmittgen, 2001). The scores of each target were normalized to those of GAPDH or β-actin. RT-PCR was conducted to confirm the expression of *retinoic acid receptor beta* (*RARβ*). The RT-PCR was performed using KOD One^®^ PCR Master Mix (Toyobo Co. Ltd.) under the following conditions: 35 cycles of 98°C for 10 s, 64°C for 5 s, and 68°C for 1 s. To amplify the *RARβ* fragments, two primers were designed according to a previous study (Thiede et al., 2014): forward, 5’-GTGTCAGTGCTTGTGAGGGA-3’ and reverse, 5’- TGCAGTACTGGCAGCGATTT-3’. The cDNA obtained from the negative controls (described below) was used as a template.

### Cell culture and Stimulation test

Male PGCs sorted from the gonads of each E6.5 embryo were cultured in the culture medium described by (Ezaki et al., 2020), with some modifications. Briefly, KnockOut DMEM (Thermo Fisher Scientific) was supplemented with 1% chicken serum (Thermo Fisher Scientific), 1× B-27 Supplement Minus Vitamin A (Thermo Fisher Scientific), 2 mM GlutaMAX™ (Thermo Fisher Scientific), 1× EmbryoMAX nucleosides (Merck, Darmstadt, Germany), 1× MEM Non-Essential Amino Acids Solution (Thermo Fisher Scientific), 1× antibiotic-antimycotic mixed stock solution (Nacalai Tesque, Kyoto, Japan), 0.5 mM monothioglycerol (FUJIFILM Wako Pure Chemical Co., Osaka, Japan), 1× sodium pyruvate, 10 ng/mL human FGF2 (PeproTech Inc., Rocky Hill, NJ, USA), 1 unit/mL heparin (Merck), 0.2 μM H1152 (FUJIFILM Wako Pure Chemical Co.,), and 0.2 μM Blebbistatin (FUJIFILM Wako Pure Chemical Co.,). The sorted PGCs were cultured at 38°C with 5% CO2 and 3% O2 and subcultured every 2-3 days. Cultured PGCs were observed using an inverted microscope (IX71; Olympus) and photographed with an Olympus DP70 camera (Olympus).

Since the number of PGCs isolated from male E6.5 gonads was very low, the PGCs were cultured and proliferated for approximately two months. These were then used for stimulation tests against RA. In this stimulation test, PGCs derived from five individuals and cultured independently were used (n = 5). Male PGCs, which were seeded at 2.5×10^4^ cells in a 35 mm dish and cultured for 24 h, were stimulated with all-trans-Retinoic acid (R2625; Merck) at a dose of 1 or 10 μM for 24 h. To prepare the RA-containing PGC medium, the RA powder was first dissolved in 99.5% EtOH at a dose of 5 mM. The RA solution was then diluted with culture medium. As a negative control, the cultured PGCs were stimulated with EtOH.

### Statistical Analysis

Statistical analyses for the RT-qPCR experiments were performed using Dunnett’s test to evaluate the differences between the negative control and RA-induced samples using R software (ver. 3.6.3). *P* < 0.05 was considered as statistically significant.

## Results

### Purification of PGCs using FACS

FACS was used to sort male and female PGCs from the blood of E2.5 embryos and gonads of E4.5 and E6.5 embryos, respectively (Fig. 1A). In this study, a monoclonal antibody against SSEA-1, a cell surface marker of chicken PGCs during early development (Karagenç et al., 1996; Mozdziak et al., 2006; Motono et al., 2008) was used to harvest PGCs. Fig 1B shows the cell sorting patterns obtained for each sample. After sorting, the purity of PGCs was evaluated by immunofluorescence staining using a monoclonal antibody against chicken VASA homologue (CVH), a chicken pan-germ cell marker (Tsunekawa et al., 2000). Expression of CVH was observed in the cytoplasm, and the purity of CVH-positive PGCs was > 96% regardless of developmental stage and sex (Fig. 1C; Table1). RNA-seq analysis was performed using the sorted PGCs under these conditions.

**Fig. 1.**
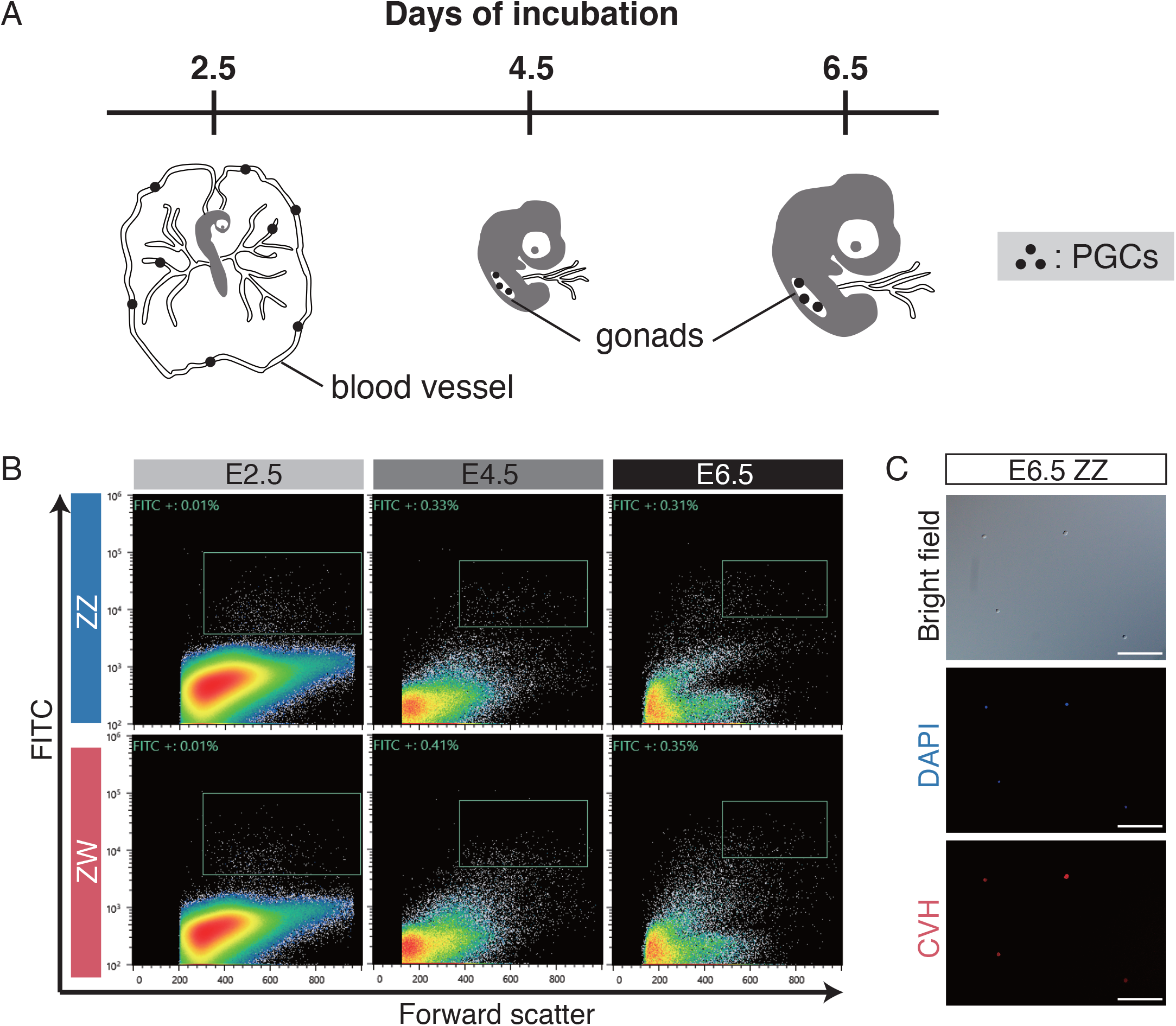
Purification of PGCs from early chick embryos. (A) A schematic illustration of localization of PGCs during embryogenesis. (B) Dot-plots of forward scatter (x-axis) versus green-fluorescence (y-axis). SSEA-1 positive cells were sorted in each stage and sex (boxes). (C) Immunofluorescence staining in sorted cells derived from E6.5 male embryos using anti-CVH monoclonal antibody. Counter staining was carried out with DAPI. Scale bars = 100 μm

**Table 1.**
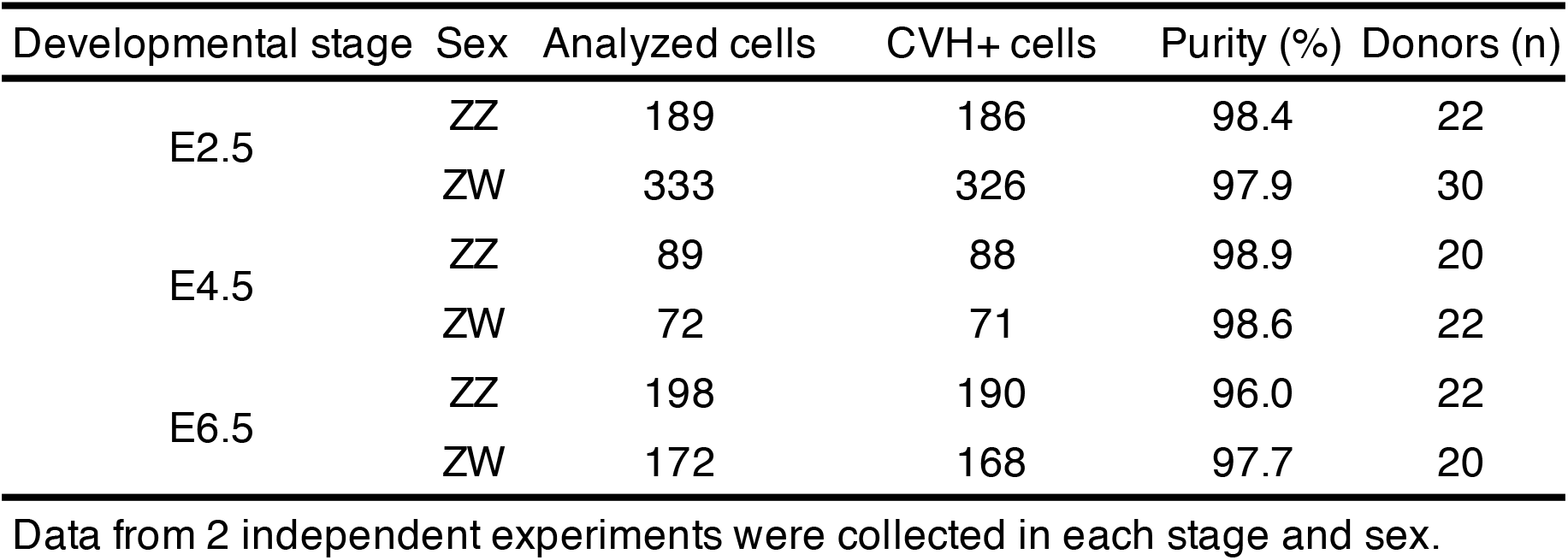
Purities of sorted PGCs in each stage and sex.

### Detection of sex-biased genes using RNA-seq analysis

To predict the sex-determination mechanism in chicken PGCs, RNA-seq analysis was performed using the sorted PGCs. Approximately, 15.7 million total reads per sample were obtained (Table S3). On average, nearly 88% of these reads were uniquely mapped to the GRCg6a reference genome.

Subsequently, we identified differentially expressed genes (DEGs) (|log2 (FC)| ≥ 3, adjusted *p*-value (Padj) < 0.05) as sex-biased genes in each developmental stage. Overall, while the number of sex-biased genes was very low in E2.5 (31 genes) and E4.5 (22 genes) embryos, it was dramatically increased in E6.5 embryos (336 genes) (Fig. 2A, Table 2). All sex-biased genes detected in this study are listed in Table S4. The number of female-biased genes was higher than that of male-biased genes at every stage. Alterations in gene expression profiles in female PGCs were observed prior to those in male PGCs.

**Fig. 2.**
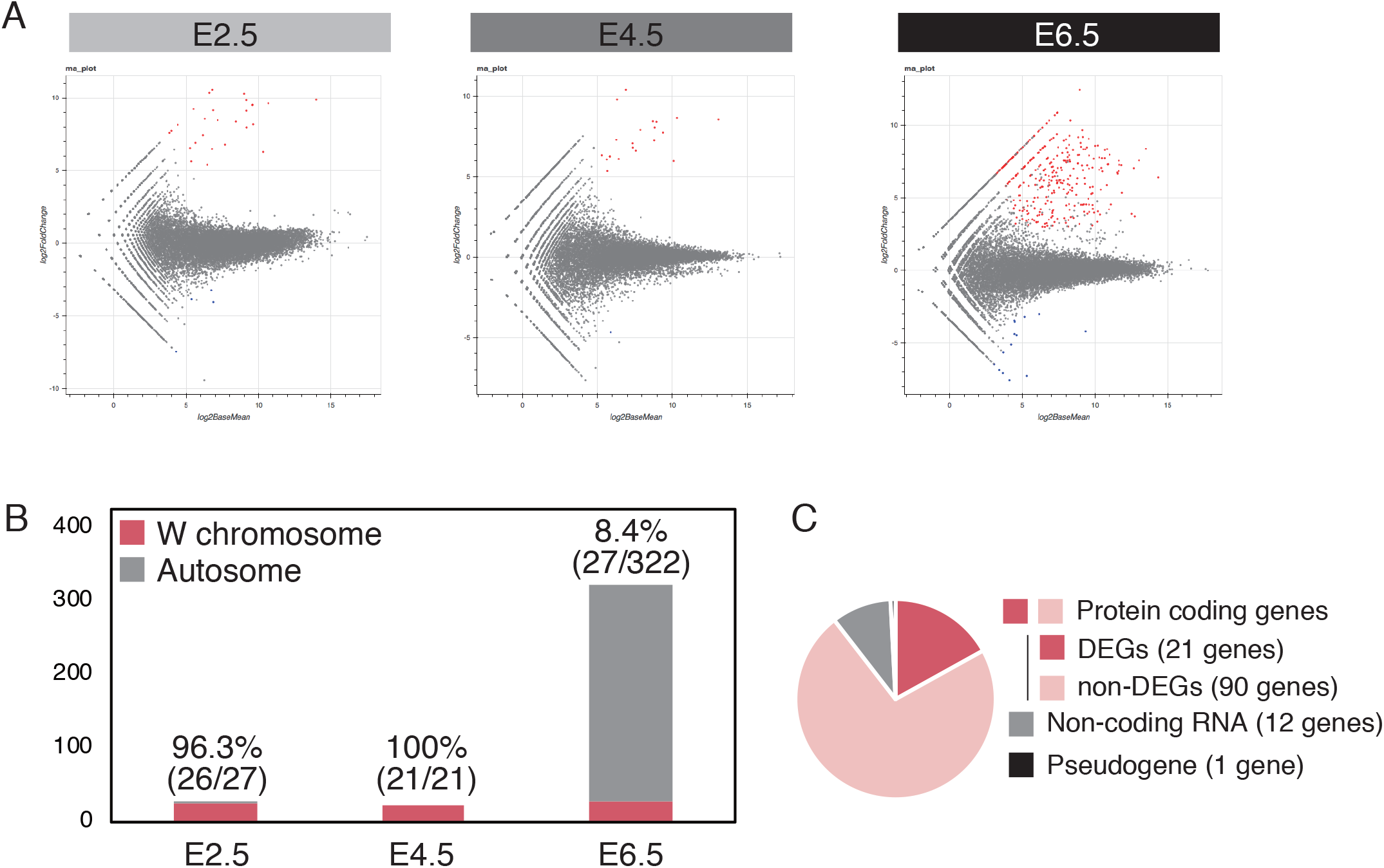
Detection and characterization of sex-biased genes during embryogenesis. (A) MA-plots of male versus female in each stage. Female-biased genes (log2 (FC) ≥ 3, Padj < 0.05) are shown in red. Male-biased genes (log2 (FC) ≤ -3, Padj < 0.05) are shown in blue. (B) Chromosomal localization of the female-biased genes in each stage. Y-axis indicates the number of genes. The ratio of W-linked gene out of whole sex-biased genes was represented on each bar. (C) Distribution of W-linked genes detected as female-biased genes in common to each stage.

**Table 2.** Number of sex-biased genes.

| Developmental stage | Male biased genes | Female-biased genes | Total sex-biased genes |
| --- | --- | --- | --- |
| E2.5 | 4 | 27 | 31 |
| E4.5 | 1 | 21 | 22 |
| E6.5 | 14 | 322 | 336 |

Chromosomal localization of female-biased genes was also analyzed at each stage. The vast majority of female-biased genes in both E2.5 and E4.5 embryos were localized in the W chromosome (Fig. 2B). In contrast, the number of female-biased genes were dramatically increased in E6.5 embryos, and 91.6% of these genes (295/322) were autosomal. Twenty-eight genes were investigated as W chromosomal female-biased genes in E2.5, E4.5, and E6.5, and 21 genes were commonly detected. All of the common female-biased genes were protein-coding genes, and corresponded to 23.3% of the W chromosome-linked protein-coding genes (Fig. 2C). The detected common female-biased genes are listed in Table 3 with functional domains predicted by Pfam, a database of protein families (Mistry et al., 2021). Gene names were annotated in only four of the 21 genes: *TGF-beta signal pathway antagonist Smad7B* (*SMAD7B*), *histidine triad nucleotide binding protein W* (*HINTW*), *heterogeneous nuclear ribonucleoprotein K-like* (*HNRNPKL*), and *ubiquitin associated protein 2* (*UBAP2*).

**Table 3.**
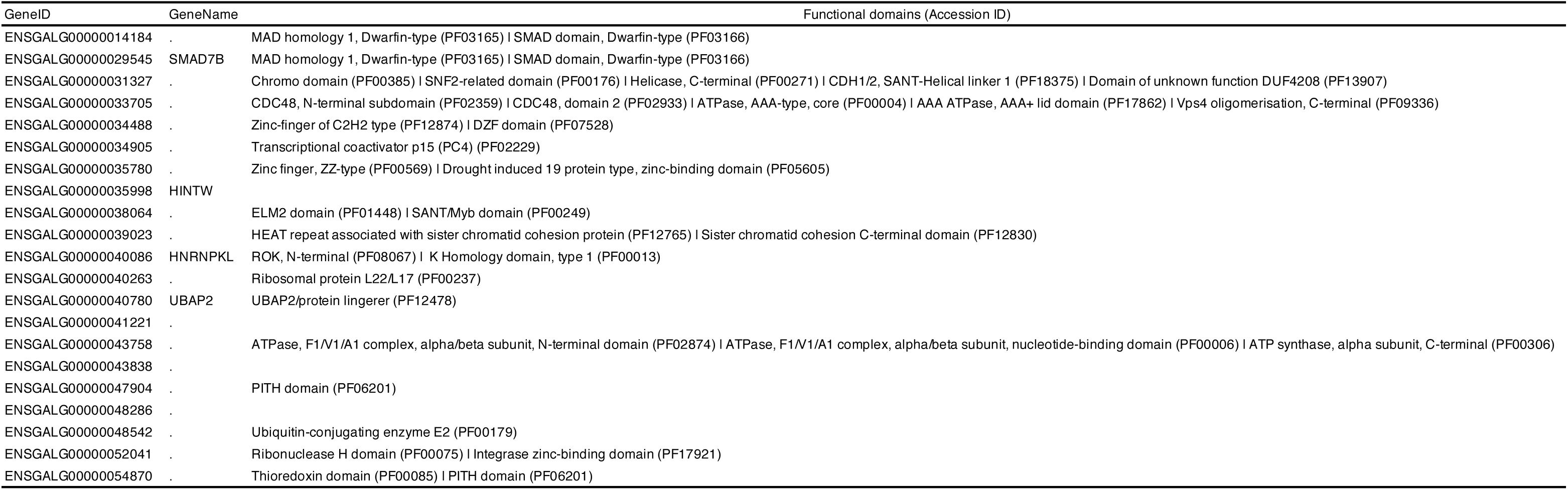
Functional information of the W-linked female-biased genes.

Functional prediction showed that several genes possess a DNA-binding domain. These were ENSGALG00000014184 (PF03165), *SMAD7B* (PF03165), ENSGALG00000034905 (PF02229), ENSGALG00000035780 (PF00569), and ENSGALG00000038064 (PF01448 and PF00249). Additionally, a chromatin remodeling-related domain (PF00385) was detected in ENSGALG00000031327. Overall, several W-linked genes were expressed and maintained in PGCs in a cell-autonomous manner, and some of them possessed functional domains that could potentially regulate transcription mechanisms.

### Enrichment analysis of female-biased genes in gPGCs derived from E6.5 embryos

Enrichment analysis was performed to characterize the features of female-biased genes detected in E6.5. First, we performed Gene Ontology (GO) analysis. Fig. 3A shows highly enriched GO terms, which are classified as biological process (BP), molecular function (MF), or cellular component (CC). Some metabolic processes were highly enriched in the BPs. In particular, organic acid metabolic processes, oxoacid metabolic processes, carboxylic acid metabolic processes, and small molecule metabolic processes were detected as major terms in female-biased genes in E6.5. In contrast, the enriched MFs were related to transporter or symporter activity, and the CCs were extracellular or apical parts. We then performed Kyoto Encyclopedia of Genes and Genomes (KEGG) pathway analysis. KEGG pathway analysis also showed enrichment of some metabolic pathways (Fig. 3B). Finally, tissue - or cell-specificity of female-biased genes was also characterized using the PaGenbase database (Pan et al., 2013). Interestingly, many female-biased genes were annotated as specific genes in the liver, kidney, and liver-derived cell lines (HEPG2 and huh-7) (Fig. 3C). All detected terms and enriched genes in each term are listed in Table S5.

**Fig. 3.**
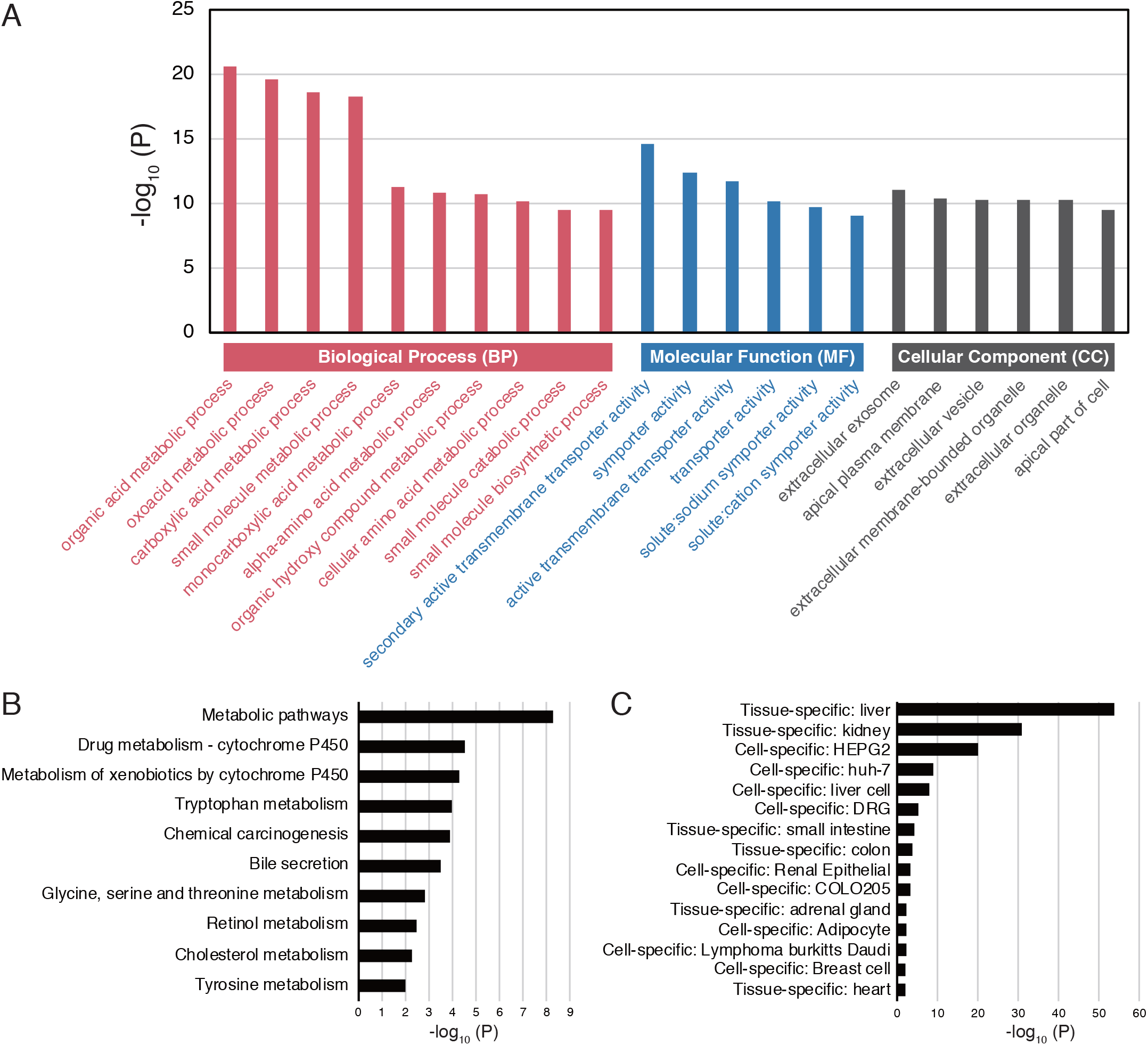
Classification of female-biased genes in E6.5 embryos. (A) The most enriched terms observed by gene ontology (GO) analysis. Three main categories, biological process (BP), molecular function (MF), and cellular component (CC), were colored by red, blue, and black, respectively. (B) The results of Kyoto Encyclopedia of Genes and Genomes (KEGG) pathway analysis. (C) The results of tissue- or cell-specificity.

To reveal the protein-protein interaction (PPI) network of female-biased genes in E6.5, we further analyzed the STRING network (Fig.4). Subsequently, 204 and 326 nodes and edges were obtained, respectively, and the average node degree was 3.2. The average local clustering coefficient was 0.315. The PPI enrichment p-value was lower than 1.0e- 16, and this network thus significantly interacted rather than being selected at random.

**Fig. 4.**
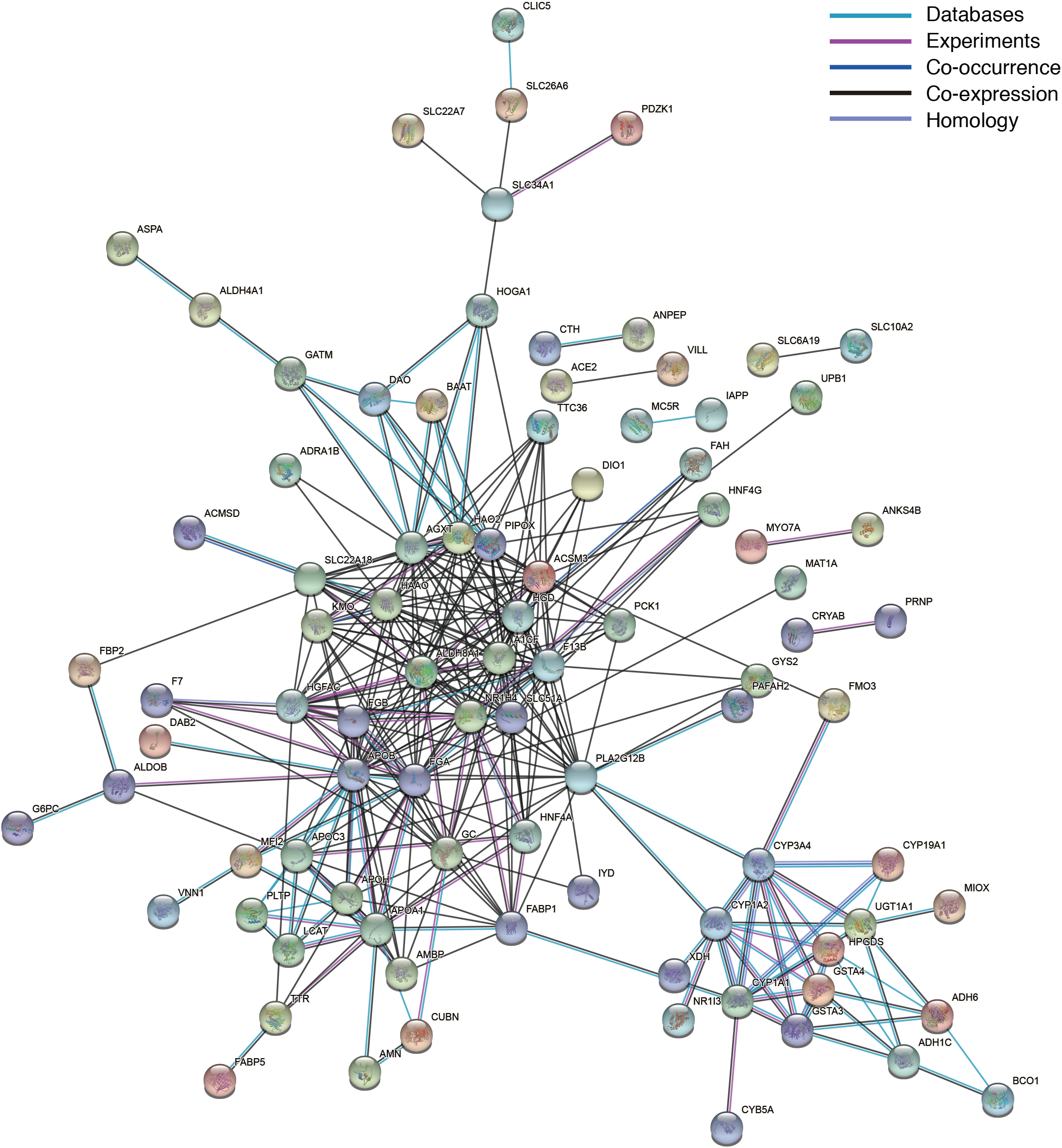
The protein-protein interaction network of female-biased genes in E6.5 chicken embryos. Nodes and edges represent protein and interaction, respectively. Different colors of edges are corresponding to different associations.

### Confirmation of the RNA-seq results using RT-qPCR

We examined the expression levels of several genes, including regucalcin (RGN), cytochrome b5 type A (CYB5A), UDP glucuronosyltransferase family 1 member A1 (UGT1A1), apolipoprotein A1 (APOA1), hematopoietic prostaglandin D synthase (HPGDS), glycine amidinotransferase (GATM), aldolase, fructose-bisphosphate B (ALDOB), and beta-ureidopropionase 1 (UPB1). These genes were included in all four major GO terms in the GO analysis: organic acid metabolic process, oxoacid metabolic process, carboxylic acid metabolic process, and small molecule metabolic process.

Moreover, *RGN*, *CYB5A*, *UGT1A1*, *APOA1*, *ALDOB*, and *UPB1* were also enriched as liver-characteristic genes. Although *APOA1* and *ALDOB* were also highly expressed in gPGCs derived from E4.5 female embryos, high expression levels of all genes were commonly observed in those from E6.5 female embryos (Fig. 5). The results of RT-qPCR corresponded to those in RNA-seq in terms of up-regulation of the candidate genes in E6.5 female gPGCs.

**Fig. 5.**
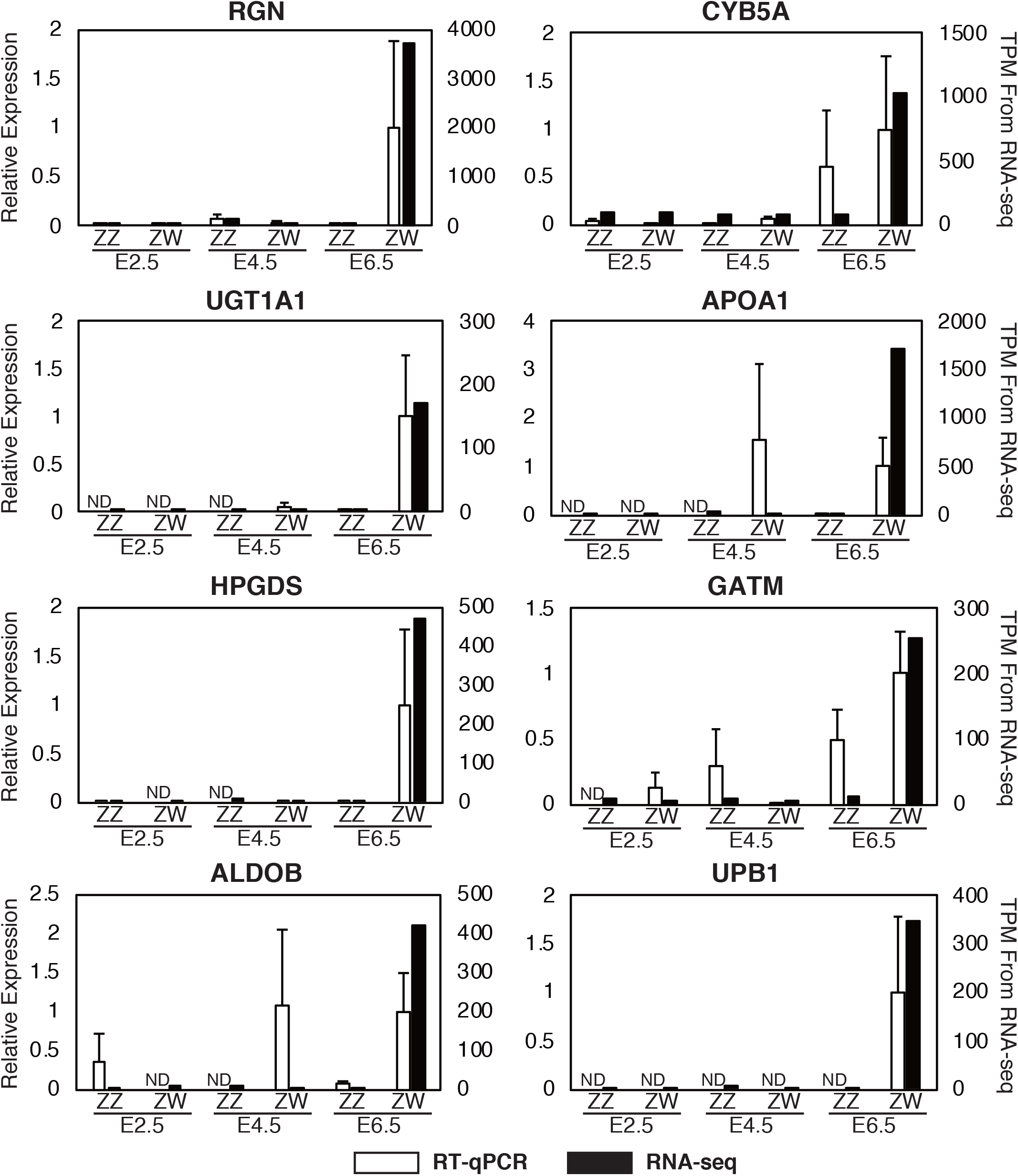
Expression analysis of the female-biased genes detected in E6.5 chicken embryos using RT-qPCR. The left y-axis indicates the score of RT-qPCR. The right x-axis indicates the scores of TPM. Open and closed bars indicate the results of RT-qPCR and RNA-seq, respectively. The 2^-ΔΔCt^ method was used for the calculation of relative expression levels, which were normalized by levels of *GAPDH*. Error bars indicate SE of triplicates (n = 3). ND means not detected.

### Retinoic acid stimulation of gPGCs in vitro

To investigate the factors involved in the upregulation of female-biased genes in gPGCs derived from E6.5 female embryos, a stimulation test using cultured gPGCs was performed. We focused on retinoic acid (RA) as a candidate factor involved in the upregulation of these genes. RA directly induces meiosis in female germ cells during embryogenesis, and thus RA is a significant factor in the feminization of PGCs. Here, gPGCs derived from E6.5 male embryos were cultured and proliferated (Fig. 6A).

**Fig. 6.**
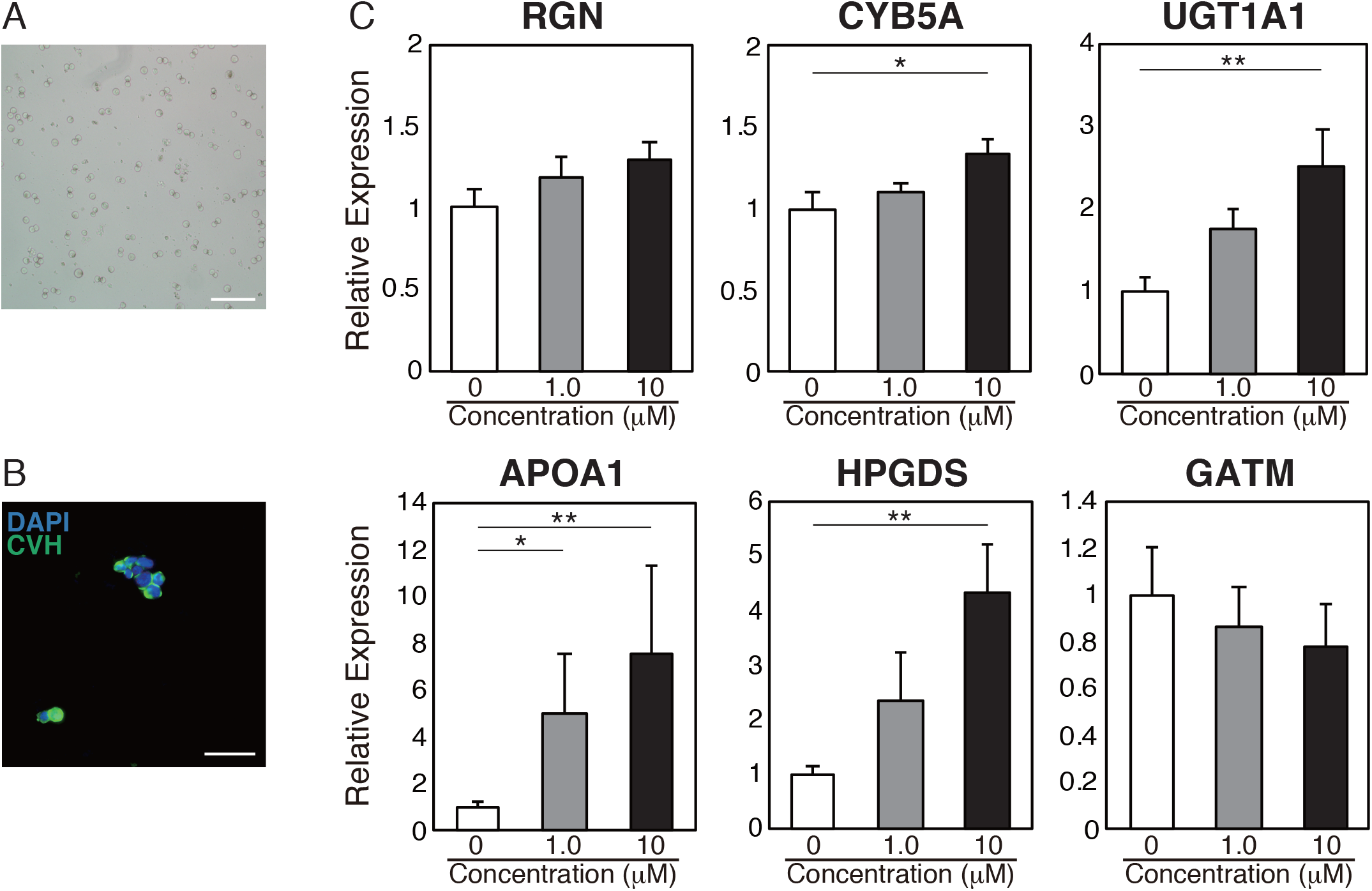
Stimulation test using RA. (A) The cultured and proliferated gPGCs derived from an E6.5 male chicken embryos. Scale bar = 100 um (B) Immunofluorescence staining using anti-CVH antibody. Counter staining was carried out with DAPI. Scale bar = 50 μm (C) Expression analysis of the female-biased genes under RA stimulation. The x-axis indicates the concentration of RA. The 2^-ΔΔCt^ method was used for the calculation of relative expression levels, which were normalized by levels of *β-actin*. Error bars indicate SE of the mean of relative expression levels during independently cultured gPGCs derived from five individuals (n = 5). Significance was evaluated by Dunnett’s test. Asterisks represent significant differences (\**P* < 0.05, \*\**P* < 0.01).

Immunofluorescence staining showed that the cultured gPGCs expressed CVH (Fig. 6B). Moreover, the expression of SSEA-1 was maintained in cultured gPGCs (data not shown). A stimulation test using RA was then conducted. The results of the expression analysis of the female-biased genes using RT-qPCR are shown in Fig. 6C. Several female-biased genes, *CYB5A*, *UGT1A1*, *APOA1*, and *HPGDS*, were upregulated in RA in a dose-dependent manner. Although no significance was detected, the expression of *RGN* was also induced. In contrast, the expression of *GATM* was decreased rather than increased, but no significant difference was detected. *ALDOB* and *UPB1* were not detected in this test, regardless of RA stimulation. Additionally, the expression of *retinoic acid receptor beta* (*RARβ*), an RA receptor, is prominently expressed in male and female embryonic gonads (Smith et al., 2008) and in PGCs (Fig. S1A), was maintained in cultured PGCs without RA stimulation (Fig. S1B). Overall, RA can enhance the upregulation of several female-biased genes in male-derived gPGCs via RARβ. Furthermore, other factors, including cell-autonomous differentiation, might be involved in the feminization of gPGCs.

## Discussion

The main goal of this study was to predict the sex-determination mechanism in avian PGCs using RNA-seq analysis. Therefore, female-specific alterations in gene expression profiles were investigated and identified. Subsequently, it was observed that sex may be established in females earlier than in males. To the best of our knowledge, this is the first study to reveal the dynamics of gene expression during sex determination of avian PGCs.

### The difference in the number of sex-biased genes between male and female

In the present study, we purified PGCs from embryonic blood and gonads using FACS (Fig. 1; Table 1), and gene expression profiles were revealed by RNA-seq analysis. The number of female-biased genes was higher than that of male-biased genes during early development (Fig. 2 and Table 2). Thus, this study sheds light on the female-specific sex-differentiation mechanism in terms of gene expression profiles.

Interestingly, avian gonadal sex determination has the opposite effect. *DMRT1*, a candidate gene in avian gonadal sex determination on the Z chromosome, is expressed in both sexes from approximately E4.5. In birds, sex-chromosome dosage compensation is incomplete (McQueen and Clinton, 2009), and the expression levels of *DMRT1* in males are also higher than in females because of the Z-linked gene (Smith et al., 1999). The strong expression of *DMRT1* induces the upregulation of masculinization-related genes, such as *HEMGN* and *SOX9*, and testicular development proceeds (Lambeth et al., 2014). In contrast, low expression levels of *DMRT1* in females result in ovarian development, including upregulation of *FOXL2* (Smith et al., 2009a; Ioannidis et al., 2021), a transcription factor that prevents testicular development (Major et al., 2019). Therefore, gonadal sex determination is likely triggered by male-specific gene expression. Indeed, the gene expression profiles of chick gonads in E4.5 showed that the number of male-biased genes was higher than that of female-biased genes (Ayers et al., 2013). Additionally, Ayers and colleagues also revealed that the increase of male-biased genes was larger than that of female-biased genes from E4.5 to E6.0 (Ayers et al., 2015). Overall, although avian gonadal sex determination is triggered by male-specific gene expression, our present study suggests that the sex determination of avian PGCs proceeds predominantly in females.

### W-linked female-biased genes and cell-autonomous sex-determination

We detected 21 W-linked female-biased genes, which were consistently expressed from E2.5 to E6.5 (Fig. 2B, C and Table3). Since the W-linked genes were upregulated in female cPGCs, these genes could be expressed independently of signals from gonadal somatic cells.

Among the 21 W-linked genes, some candidates were identified as potential regulators in the establishment of sexual identity. ENSGALG00000014184 and ENSGALG00000029545 (*SMAD7B*) possess the SMAD domain, which is conserved in the SMAD family. The SMAD family is highly phosphorylated in chicken PGCs, and the activation of SMAD signaling is essential for the *in vitro* proliferation of PGCs (Whyte et al., 2015). The *SMAD7B* gene was cloned in a previous study (Vargesson and Laufer, 2009). However, the functions of SMAD7B have not been determined and neither has ENSGALG00000014184. Conversely, ENSGALG00000035998 (histidine triad nucleotide-binding protein W; HINTW) is a well-analyzed W-linked candidate characterized by strong expression patterns in female embryos (Hori et al., 2000; O’Neill et al., 2000). Although overexpression of HINTW did not affect gonadal masculinization (Smith et al., 2009b), the function of sex determination in PGCs remains unclear. Additionally, several candidates that potentially regulate the expression of other genes were also identified in this study. In the future, functional analysis of W-linked genes in PGCs is required.

Several previous reports also may support sexual differences in chicken PGCs at the pre-gonadal stage. First, sexual bipotency was lost in cPGCs. Compared to PGC precursor cells included in blastodermal cells, cPGCs hardly differentiate into gametes in the opposite sex gonads following transplantation (Kagami et al., 1997; Naito et al., 1999; Tagami et al., 2007). Proliferation activity of PGCs might also be different between males and females. The number of female PGCs concentrated in the intermediate mesoderm in E2.5 embryos was significantly higher than that of male PGCs (Nakamura et al., 2007). Our study provides new insights for some candidates that can contribute to the gonadal differentiation-independent sex determination of PGCs.

### Characters of female-biased genes detected in E6.5 embryos

In E6.5 embryos, female PGCs possessed a large number of sex-biased genes when compared to male PGCs (Fig. 2A). These female-biased genes were significantly enriched in some metabolic processes (Fig. 3A and 3 B) and interacted more strongly than randomly selected genes (Fig. 4). Many female-biased genes were annotated as liver- or kidney-characteristic genes (Fig. 3C). RT-qPCR analysis further confirmed the female-biased expression patterns of candidate genes, that is, *RGN*, *CYB5A*, *UGT1A1*, *APOA1*, *HPGDS*, *GATM*, *ALDOB*, and *UPB1* (Fig. 5). These results suggest that female PGCs undergo metabolic processes at E6.5, which may contribute to the differentiation of female germ cells.

After gonadal sex determination, chick germ cells showed sex-specific developmental patterns. While male germ cells barely proliferate during embryonic testicular development, female germ cells obtained high proliferating activity from at least E9.0 (Hughes, 1963; Méndez et al., 2005). Then, female germ cells undergo meiosis in E15.5. The metabolic processes of female PGCs at E6.5, may contribute to the female germ cell-specific proliferation activity.

Previous studies have shown the activation of several metabolic processes in PGCs. Prostaglandin D2 (PGD2), which is synthesized by the HPGDS and Lipocalin-type prostaglandin D synthase (LPGDS), is produced in murine testicular PGCs, and related to sex-differentiation of those (Moniot et al., 2014). Calcium (Ca^2+^) homeostasis also contributes to gametogenesis. RGN is a Ca^2+^-binding protein and is involved in the regulation of several enzymatic activities, such as cell proliferation and apoptosis.

Previously focused on as a characteristic factor of the liver and kidney (Shimokawa and Yamaguchi, 1992; Yamaguchi and Isogai, 1993), RGN is also expressed in mammalian testes, including germ cells, and is likely related to spermatogenesis (Laurentino et al., 2011; Laurentino et al., 2012). Furthermore, since murine early embryonic PGCs exhibit high glycolytic activity (Hayashi et al., 2017), the contribution of the glycolytic pathway, including activity of the ALDOB, to the development of these pathways is suggested. Additionally, gene profiling analyses comparing chick gPGCs and blastoderms under mixed-sex conditions showed upregulation of *APOA1*, *RGN*, and nucleotide metabolism-related genes, including *UPB1*, in the gPGCs (Rengaraj et al., 2013; Rengaraj et al., 2014). Our study provides new insights that the activation of these metabolic process-related genes is a characteristic feature of female gPGCs, at least in the early developmental stage.

### Influence of Retinoic acid against sex-determination in embryonic germ cells

Stimulation of male PGCs obtained from the gonads of E6.5 embryos with RA *in vitro* resulted in the upregulation of several female-biased genes (Fig. 6). This suggests the contribution of somatic cell-derived signals, including RA, to the feminization of gPGCs at E6.5. RA-independent regulatory factors could also be involved in the feminization of chicken PGCs because RA had no effect on the expression of *GATM* and *RGN*.

RA directly induces meiosis, which is a female-specific developmental pattern during embryogenesis (Bowles et al., 2006; Koubova et al., 2006). In developing chicken gonads, RA is mainly synthesized by retinaldehyde dehydrogenase 2 (RALDH2) from E6.5, throughout development (Smith et al., 2008). On the other hand, a RA-degrading enzyme, cytochrome P450 family 26, subfamily B member 1 (CYP26B1), is also expressed in the gonads at the same time (Smith et al., 2008). The expression of CYP26B1 in female gonads is decreased during development, and this results in meiosis at E15.5 via upregulation of stimulated by retinoic acid 8 (STRA8), a premeiotic marker (Oulad-Abdelghani et al., 1996; Baltus et al., 2006), by RA stimulation from E12.5. In contrast, the expression of CYB26B1 increases up to E9.5 and maintains its expression levels during embryogenesis in male gonads (Smith et al., 2008). Our findings suggest that RA also regulates the upregulation of several female-biased genes in female PGCs at E6.5, prior to STRA8 expression. To confirm this hypothesis, further studies are needed to verify the expression of RA and analyze RA functions during gonadal sex determination *in vivo*.

As an RA-independent regulatory factor, sexual differences in DNA methylation patterns were predicted. A previous study demonstrated that the methylation in *GATM*gene is significantly higher in E6.0 male gPGCs compared to that in female gPGCs (Jang et al., 2013). This is consistent with our findings that RA stimulation could not induce *GATM* expression in male gPGCs obtained from embryos at E6.5. Additionally, this also suggests cell-autonomous sex differentiation in PGCs after gonadal sex determination.

## Conclusion

In this study, we examined the gene expression profiles of chick PGCs during gonadal sex determination. Alterations in gene expression profiles were observed in females prior to that in males. The feminization of chick PGCs might be induced by RA as well as other factors, including a cell-autonomous mechanism. Thus, our findings elucidate the mechanism of avian-specific sex-determination and reveal valuable insights into vertebrate evolution associated with this mechanism.

However, the present study has only projected the underlying molecular mechanisms of feminization in chick PGCs. Therefore, further investigations are warranted to conclusively establish the functional correlations between the female-biased genes and the sex-determination of PGCs. Since 2014, an effective strategy to produce genetically modified chickens using genome editing tools is being extensively used (Park et al., 2014). This protocol is also used for basic research of germ cell development (Taylor et al., 2017). Further studies and functional analyses of genetic modifications may identify the key molecule(s), which induce(s) feminization of chick PGCs, among the female-biased genes.

## Acknowledgment

We would like to thank K.K. DNAFORM (Yokohama, Japan) for the RNA-seq analyses. We thank Editage (http:// www.editage.jp) for English language editing.

## Data availability

All RNA-seq data sets are publicly available at Gene Expression Omnibus under accession number GSE188689.

