## Supplementary material for "Prediction of sex-determination mechanisms in avian primordial germ cells using RNA-seq analysis": Suppl_figure

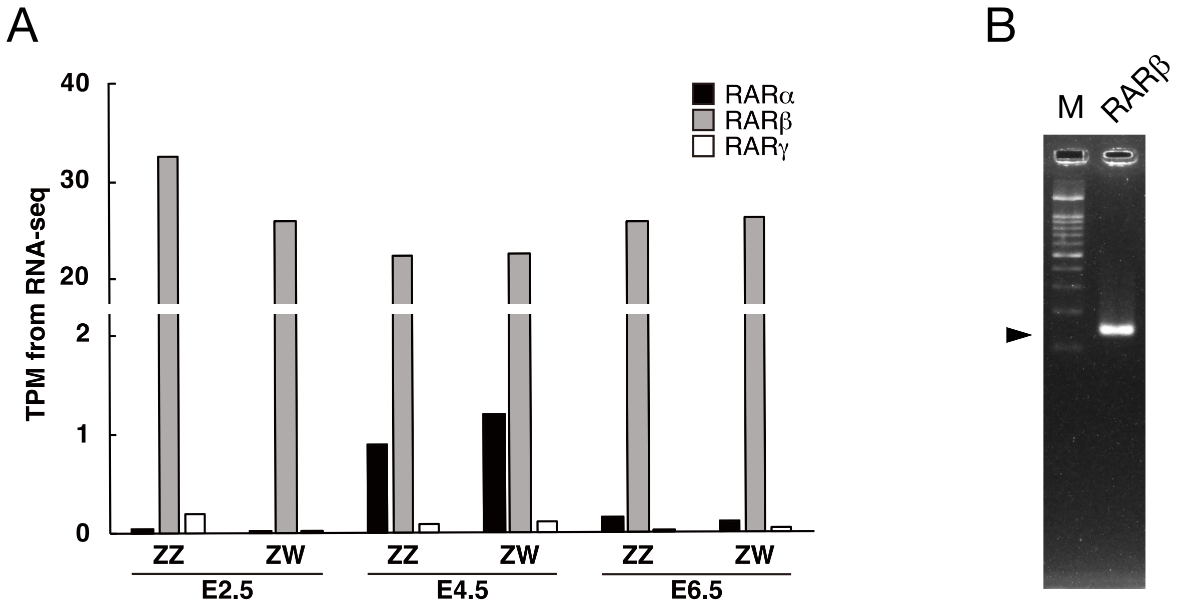


Fig. S1. Evaluation of the expressions of Retinoic acid receptors (RARs). (A) The expression levels of all RARs, RARα, RARβ, and RARγ, determined by RNA-seq analysis in TPM units in each sex and developmental stage. (B) Gel electrophoresis of the RT-PCR fragment. The predicted size of the fragment is 129 bp (indicated by arrowhead). M, 100 bp marker.
